# iScore: A novel graph kernel-based function for scoring protein-protein docking models

**DOI:** 10.1101/498584

**Authors:** Cunliang Geng, Yong Jung, Nicolas Renaud, Vasant Honavar, Alexandre M.J.J. Bonvin, Li C. Xue

## Abstract

Protein complexes play a central role in many aspects of biological function. Knowledge of the three-dimensional (3D) structures of protein complexes is critical for gaining insights into the structural basis of interactions and their roles in the biomolecular pathways that orchestrate key cellular processes. Because of the expense and effort associated with experimental determination of 3D structures of protein complexes, computational docking has evolved as a valuable tool to predict the 3D structures of biomolecular complexes. Despite recent progress, reliably distinguishing near-native docking conformations from a large number of candidate conformations, the so-called scoring problem, remains a major challenge. Here we present iScore, a novel approach to scoring docked conformations that combines HADDOCK energy terms with a score obtained using a graph representation of the protein-protein interfaces and a measure of evolutionary conservation. It achieves a scoring performance competitive with, or superior to that of the state-of-the-art scoring functions on independent data sets consisting docking software-specific data sets and the CAPRI score set built from a wide variety of docking approaches. iScore ranks among the top scoring approaches on the CAPRI score set (13 targets) when compared with the 37 scoring groups in CAPRI. The results demonstrate the utility of combining evolutionary and topological, and physicochemical information for scoring docked conformations. This work represents the first successful demonstration of graph kernel to protein interfaces for effective discrimination of near-native and non-native conformations of protein complexes. It paves the way for the further development of computational methods for predicting the structure of protein complexes.

## INTRODUCTION

Protein-protein interactions (PPIs) play a crucial role in most cellular processes and activities such as signal transduction, immune response, enzyme catalysis, etc. Getting insight into the three dimensional (3D) structures of those protein-protein complexes is fundamental to understand their functions and mechanisms^1,2^. Due to the prohibitive cost and effort involved in experimental determination of the structure of protein complexes^3^, computational modelling, and in particular docking, has established itself as a valuable complementary approach to obtaining insights into structural basis of protein interactions, interfaces, and complexes^4–10^.

Computational docking typically involves two steps^4,7–9^: Sampling, i.e., the search of the interaction space between two molecules to generate as many as possible near-native models; and scoring, i.e., the identification of near-native models out of the pool of sampled conformations. As shown in the community-wide Critical Assessment of PRediction of Interactions (CAPRI)^11–14^, scoring is still a major challenge in the field. There is thus still plenty of room to improve the scoring functions used in protein-protein docking^10,15^.

Scoring functions can be classified into three types: *i)* physical energy term-based, *ii)* statistical potential-based and *iii)* machine learning-based. Physical energy-based scoring functions are usually a weighted linear combination of multiple energetic terms. These are widely used in many docking programs such as HADDOCK^16,17^, SwarmDock^18^, pyDock^19–21^, ZDock^22,23^, and ATTRACT^24^. Taking HADDOCK as an example, its scoring function consists of intermolecular electrostatic and van der Waals energy terms combined with an empirical desolvation potential^25^ as well as a buried surface area (BSA)-based term depending on the stage of the protocol^17^. Statistical potential-based scoring functions such as 3D-Dock^26^, DFIRE^27^, and SIPPER^28^, typically convert distance-dependent pairwise atom-atom or residue-residue contacts distributions into potentials through Boltzmann inversion. Unlike classical scoring functions that consist of linear combinations of energy terms, or simple geometric and physicochemical features^29–31^, a machine learning approach can discover complex nonlinear combinations of features of protein-protein interfaces to train a classifier to label a docking model as near-native model or not. Simple machine learning algorithms work with fixed dimensional feature vectors. Because interfaces of different docking models can vary widely in size and shape, and in the arrangement of their interfacial residues, most machine learning based scoring functions typically use global features of the entire interface, for example, the total interaction energy and the BSA. However, such an approach fails to effectively utilize details of the spatial arrangement of interfacial residues/atoms.

Graphs, in which the nodes encode the amino acid residues or atoms and the intermolecular contacts between them are encoded by the edges, offer a natural and information-rich representation of protein-protein interfaces. Unlike the global interface feature vectors described above, a graph has a residue- or atom-level resolution and naturally encodes the topological information of interacting residues/atoms^32,33^. Furthermore, the size of a graph is not fixed and can vary depending on the size of the interface.

Such graph-based descriptions have been used previously in several scoring functions^34–36^. Graph (or network) topology-based metrics have mostly been used. Chang et al. 2008^34^ exploited node degrees (measuring the number of direct contacts of a node) and clustering coefficients (measuring how likely a node and its neighbours tend to form a clique) to score docking models. Similarly, Pons et al. 2011^35^ used closeness (measuring how far a node from the rest of the nodes in a network) and betweenness (measuring how important a node as a connector in a network) in scoring with the intuition that residues with high centralities in a network tend to be key functional residues. Unlike the network topology-based approaches, the SPIDER^36^ scoring function uses a graph to represent the interface at residue level with nodes labelled by their amino acid identity. It ranks the docking models by counting the frequency of native motifs in the interface graph. However, all the preceding fail to fully exploit the rich features of protein interfaces.

Against this background, we represent the interface with a labelled graph, where the nodes encode the interface residues, edges encode residue-residue contacts, and the nodes are annotated with evolutionary conservation profiles. We treat the scoring problem as a binary classification problem. By calculating the similarity between an interface graph from a docking model with the positive (native) and negative (non-native) interface graphs in the training set, we predict the likelihood of the query interface graph belonging to the positive class or the negative class (Figure 1). We make use of a novel *graph kernel* to compute the pair-wise similarity between the graph representations of protein-protein interfaces. We call the resulting graph kernel-based scoring function GraphRank.

**Figure 1.**
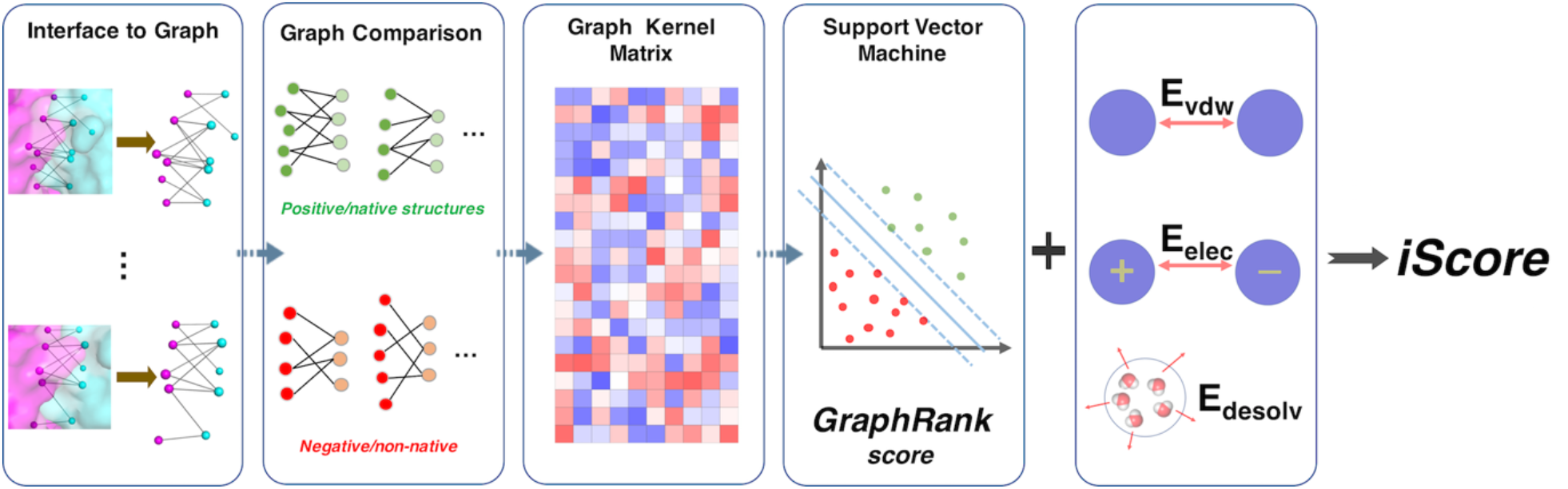
Schematic workflow of our graph kernel-based scoring method. Docking models for a protein-protein complex are first represented as graphs by treating the interface residues as graph nodes and the intermolecular contacts they form as graph edges. Interface features are added to the graph as node or edge labels (only PSSM profiles as node labels in this case). Then, each of the interface graphs of the docking models is compared to the interface graphs of both the positive (native) structure and negative (non-native) models. This graph comparison generates a similarity matrix for the docking models with the number of rows and columns corresponding to the number of docking models and the total number of positive and negative graphs, respectively. Next, the support vector machine takes the graph kernel matrix as input and predicts decision values that are used as the GraphRank score. The final scoring function iScore is a linear combination of the GraphRank score and HADDOCK energetic terms (van der Waals, electrostatic and desolvation energies). The weights of this linear combination are optimized using the genetic algorithm (GA) over the BM4 HADDOCK dataset.

GraphRank exploits random walk graph kernel (RWGK)^37^ for computing the similarity of labeled graphs, which has previously been used for protein function prediction^38^ to calculate the similarity between two interface graphs. By simultaneously conducting random walks on two graphs, RWGK measures the similarity of two graphs by aggregating the similarity of the set of random walks on the two graphs. Unlike previous graph-based scoring functions, RWGK allows GraphRank to fully exploit various node labels and edge labels and to explicitly specify the starting and ending probability of the random walks. GraphRank has two major advantages over classical machine learning based scoring functions. First, GraphRank uses a more detailed representation of protein interfaces than that provided by the fixed dimensional feature vectors used by classical machine learning approaches. GraphRank exploits residue level attributes and network topology. Second, GraphRank uses the full profile of interface conservation as node labels, i.e., each node is represented as a 20 by 1 vector of conservation profile extracted from the Position Specific Scoring Matrix (PSSM). Residue conservation information plays an important role in protein-protein recognitions^39–41^ and hence different types of conservation information have been used in several existing scoring functions^42–44^. The PSSM is a multiple-sequence-alignment (MSA) based conservation matrix. Its value is a log likelihood ratio between the observed probability of one type of amino acid appearing in a specific position in the MSA and the expected probability of that amino acid type appearing in a random sequence. Each position in a protein can be represented as a 20 by 1 PSSM profile, which captures the conservation characteristic of each amino acid type at a specific position.

For GraphRank we designed a specific random walk graph kernel to compare interface graphs. A graph similarity matrix was calculated from a balanced dataset of native and non-native structures from the protein-protein docking benchmark version 4.0 (BM4), and was used to train a support vector machine (SVM) classifier. GraphRank, the resulting scoring function, uses only the residue conservation information as node labels and as the basis of starting and ending probabilities of random walks. We further combined the GraphRank score with intermolecular energies, resulting our final scoring function, iScore. We benchmarked the iScore and GraphRank scoring functions on two independent sets of docking models for two different purposes: 1) 4 sets of *docking software-specific* models and their respective scoring functions and 2) the CAPRI score set, a set of *docking software-nonspecific* models, in which models from different docking programs are mixed together. The results of our experiments on both benchmarks show that iScore achieves scoring performance that is competitive with or superior to that of the state-of-the-art scoring functions. These results represent the first successful demonstration of the use of graph kernel applied to protein interfaces for effective discrimination of near-native and non-native conformations of protein complexes.

## METHODS

### Constructing interface graph and random walk graph kernel

#### Representing protein-protein interfaces as labelled bipartite graphs

A residue is defined as an interface residue if any of its atoms is within 6Å of any atom of another residue in the partner protein. This is a commonly used interface definition^45^, and, for example, a similar cutoff (5.5Å) has been shown to work well for contacts-based binding affinity prediction^46^. We represent the interface of a native complex or a docking model as a bipartite graph (Figure 1), in which each node is an interface residue, and each edge consists of two nodes that are within 6Å distance from each other (based on any atom-atom distance within 6Å between those residues). We further label the graph node with residue conservation profiles from Position Specific Scoring Matrix (PSSM). Each node is thus represented by a 20×1 vector of PSSM profile. Our current implementation uses a single type of nodes, namely residues, labeled with their PSSM profiles, and a single type of edges, namely, those that encode inter-residue contacts. However, our framework admits multiple types of nodes and edge labels.

The PSSM was calculated through PSI-BLAST^47^ of BLAST 2.7.1+. The parameters of the BLAST substitution matrix, word size, gap open cost and gap extend cost were automatically set based on the length of protein sequence using the recommended values in the BLAST user guide (https://www.ncbi.nlm.nih.gov/books/NBK279084/) (see **Table S1**). Other parameters were: Number of iterations set to 3 and the e-value threshold to 0.0001. The BLAST database used was the nr database (the non-redundant BLAST curated protein sequence database), version of February 04, 2018.

#### Random walk graph kernel for interface graphs

We define a random walk graph kernel (RWGK) to measure the similarity of two interface graphs. Given two labeled graphs, a RWGK first applies simultaneous random walks on the two graphs with the same walk length (the number of edges) and then calculates the similarity between those two random walks. The RWGK score is then the weighted sum of the walk similarity varying the walk length from 0 to infinity^48^.

Gärtner et al.^49^ proposed an elegant approach for calculating all random walks within two graphs using direct product graphs. A graph *G* consists of a set of *n* nodes *V* = {*v*_1_, *v*_2_, …, *v_n_*} and a set of *m* edge *E* = {*e*_1_, *e*_2_, …, *e_m_*} where the edge *e*_*i*_ is defined by two nodes. Given two graphs *G* = {*V, E*} and *G*′ = {*V*′, *E*′}, the direct product graph *G*_×_ is a graph defined as follows:

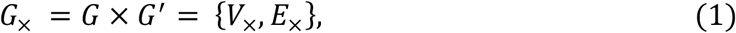

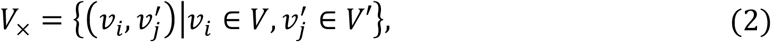

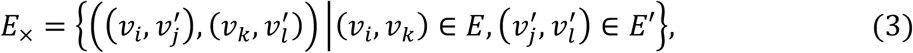

where *V*_×_ is the node set and *E*_×_ is the edge set. In other words, *G*_×_ is a graph over pairs of nodes from *G* and *G*′, and two nodes in *G*_×_ are neighbors if and only if the corresponding nodes in *G* and *G*′ are both neighbors^37^.

The simultaneous random walks on graphs *G* and *G*′ are equivalent to a random walk on the direct product graph *G*_×_. In other words, each walk on the direct product graph *G*_×_ corresponds to two walks on the two individual graphs, allowing the calculation of a similarity score between them. When the walk length is 1, these similarity scores are the elements of the weight matrix *W*_×_ of 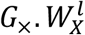 consists of similarity scores of walk length of *l*. The similarity between graphs *G* and *G*′ is thus the weighted sum of these walk similarities.

Formally, the random walk graph kernel is originally defined by Vishwanathan et al.^37^ as:

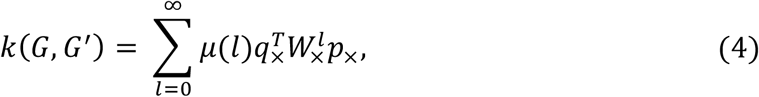

where *l* is the length of random walk on *G*_×_, *μ*(*l*) is a factor that allows one to (de-)emphasize walks with different lengths, *W*_×_ is the weight matrix of *G*_×_, and *q*_×_ and *p*_×_ are the starting and stopping probabilities of random walks on *G*_×_, respectively. In our study, we limit the maximum walk length to 3, and *μ*(*l*) is set to 1 for *l* = 0 to 3.

And *W*_×_, *q*_×_ and *p*_×_ are designed as follows.

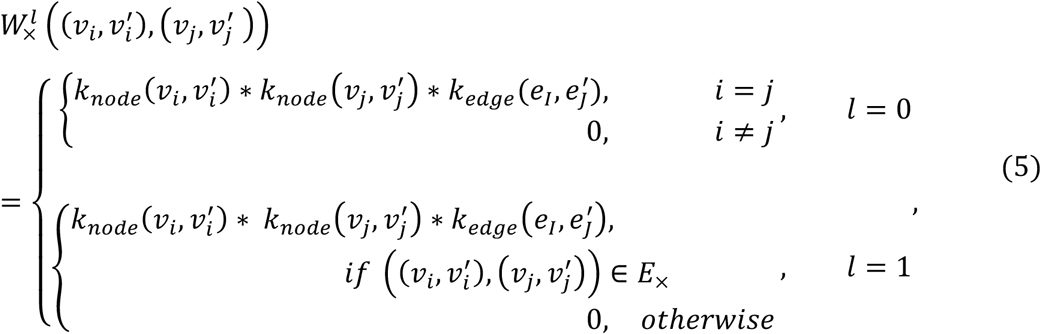

where 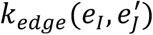 is the kernel to measure the similarity between two edges, *e_I_* = (*ν_i_, ν_j_*) and 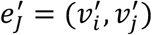. Since we do not use specific edge labels here, 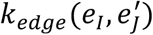 is simply set to 1. 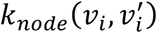 is the kernel to measure similarity between nodes defined as follows:

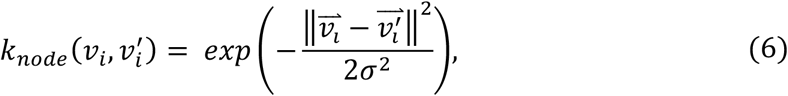

where 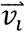 and 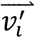 are node labels for nodes *v_i_* and 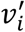, respectively. As described above, we used PSSM residue conservation profiles as node label. *σ* was set to 10 by simply checking the distribution of some 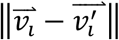 values.

We bias the random walks to start and end with conserved residues by giving those higher starting and ending probabilities. For this, we define the starting and ending probabilities 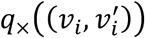 and 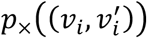 from the normalized conservation score as follows:

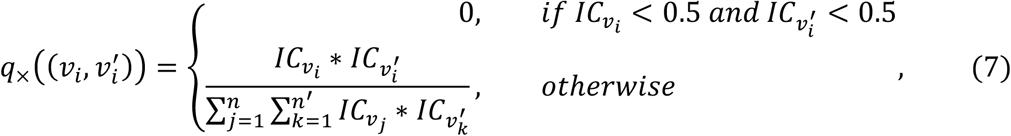

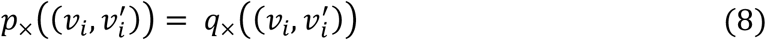

where 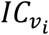 and 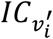 are the PSSM information content (IC) for the nodes *v_i_* and 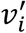, respectively, and *n* and *n*′ are the number of nodes in graph *G* and *G*′, respectively. IC is always ⩾0. The higher the IC, the more conserved a residue is.

#### Support vector machine (SVM) algorithm

SVM is a kernel-based learning algorithm^50,51^. We used the SVM implementation from the LIBSVM^52^ package to train a scoring function taking the *N* × *N* graph kernel matrix from the training dataset as input (*N* is the number of the training graphs). Given a test data (an interface graph of a docking model in our case), we calculate the kernel vector that consists of the similarities of this query graph with all the training graphs. The trained SVM-based scoring model uses the resulting vector of similarities of the query graph with all of the training graphs as well as the labels of the training graphs to predict the likelihood of the query graph corresponds to a near-native conformation.

### Evaluation Metrics to compare scoring functions

We used the success rate at cluster level to evaluate the scoring functions. We define a cluster as a hit if at least one of the top 4 models in that cluster is of acceptable or better quality. The success rate on top N clusters is defined as the number of cases (complexes) with at least one hit out of the N clusters divided by the total number of complexes considered.

The quality of the docking models was evaluated using standard CAPRI criteria based on the interface or ligand Root Mean Squared Deviations (i-RMSDs and l-RMSDs, respectively) and fraction of native contacts (Fnat) (for details refer to Figure 1 of Lensink et. al.^11^). They were classified as incorrect (i-RMSD>4Å or Fnat < 0.1), acceptable (2Å<i-RMSD≤4Å and Fnat ≥0.1), medium (1Å<i-RMSD≤2Å and Fnat ≥0.3) or high (i-RMSD≤1Å and Fnat ≥0.5) quality^11^.

### Training on docking benchmark 4 docking models

#### Training dataset for GraphRank

The dataset for training was based on protein-protein complexes from the protein-protein docking benchmark version 4.0 (BM4), considering only dimers, resulting in a set of 117 non-redundant protein-protein complexes. Docking models for those complexes had been generated previously by running HADDOCK in its *ab initio* mode using center of mass restraints^53^. The crystal structures of these 117 complexes (the “native” structures) form our positive training set. The average number of nodes and edges in the corresponding graphs for this native set are 68±25 and 119±55, respectively. To create a balanced training set, we randomly selected 117 non-native (wrong) models from the pool of HADDOCK models with i-RMSD ⩾ 10Å and number of graph nodes ⩾5 as our negative training set. The average number of nodes and edges in the non-native set are 48±14 and 70±23, respectively. In total, we thus have 234 (=117*2) structures as our training set.

#### Training dataset for iScore

For the training of iScore we selected BM4 complexes for which HADDOCK, running in ab-initio mode using center of mass restraints, generated at least one good model in the final water refinement stage. This resulted in 63 cases for which at least one docking model with acceptable or better quality was present in the final set of400 water-refined models. This dataset is denoted in the following as the BM4 HADDOCK dataset.

#### Training the graph kernel-based scoring function (GraphRank)

We applied the commonly-used SVM classifier C-SVC from LIBSVM^52^ to train our scoring function. We precomputed the random walk graph kernel matrix (234 × 234) for the training data and used it as input of the SVM classifier. The SVM outputs the predicted decision values for a test case (the decision values from libsvm is defined as 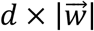, where *d* is the distance from a point to the hyperplane and 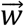 is the weight vector of SVM that defines the classification hyperplane). To be consistent with energy terms which we later incorporated into iScore (the lower the energy the better a model), we use the negative decision value from the SVM as the final score of GraphRank. The resulting optimised SVM classifier is denoted as the “GraphRank” scoring function.

#### Integrating GraphRank score with energetic terms (iScore)

We combined the GraphRank score with three energetic terms from HADDOCK to train a simple linear scoring function named iScore.

The HADDOCK energetic terms used are:

- Evdw, the intermolecular van der Waals energy described by a 12-6 Lennard-Jones potential;
- Eelec, the intermolecular electrostatic energy described by a Coulomb potential;
- Edesolv, an empirical desolvation energy term.

The van der Waals and electrostatic energies are calculated using a 8.5Å distance cutoff using the OPLS united atom force field^54^.

The GraphRank score and HADDOCK terms were normalised with the following equation:

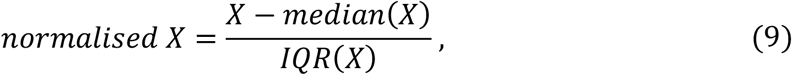

where the *X* is a set of values for a specific term, *median*(*X*) is the median value of this term, *IQR* is the interquartile range, which is the difference between the 75th and 25th percentiles.

We optimised the weights of the various iScore terms (the normalised GraphRank score and energetic features) on the BM4 HADDOCK dataset (63 cases and 400 models/case), using a genetic algorithm (GA). We used the normalised discounted cumulative gain (nDCG)^55^ to evaluate the model ranking from each combination of the GraphRank score and energetic terms. This is a common measure of ranking quality for evaluating web search engine algorithms^56^. Specifically, nDCG is defined as follows:

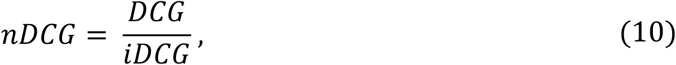

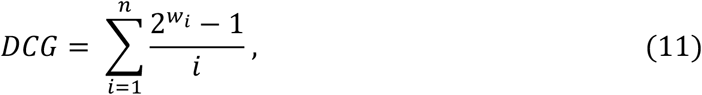

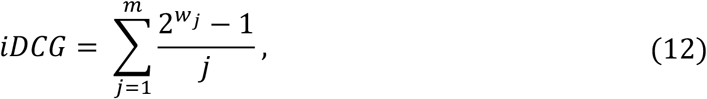

where *DCG* is the discounted cumulative gain calculated over the total number of models (here *n* in Eq. 11 is 400). *iDCG* is the ideal DCG (meaning all the hits are ranked at the top 1, 2, … m, where *m* is the total number of hits), and *nDCG* is the normalised DCG. *i* is the ranking position of a model, *w_i_* is the weight of a model ranked at position *i*. Here, we set *w_i_* = 1 if *i* is a near-native model, and *w_i_* = 0 otherwise. The contribution of a model to DCG becomes thus 0 or 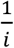, where *i* is the ranking of the model.

The fitness function for the GA optimisation was defined as the average of squared *nDCG* values for the 63 cases (one nDCG value per case). The parameters of the GA optimisation were: Population size = 800, maximum generations = 100, crossover rate = 0.8 and stopping tolerance = 0.001. The GA converged quickly, stopping at the 51th generation. The GA optimisation was repeated 30 times and the median values were used as final weights.

### Validation and comparison with state-of-the-art scoring functions

#### I. Validation on models from different docking programs

We validated iScore’s performance on docking models from four different docking programs: HADDOCK^16,57^, SwarmDock^18^, pyDock^19–21^ and ZDock^22,23^. These models were used to evaluate our scoring functions and compare them with the original scoring functions in these respective docking programs. The protein-protein complexes used for testing are the new entries from the protein-protein docking benchmark version 5.0 (BM5)^58^, on which none of the docking software listed above has been previously trained. These cases are also non-redundant to our training set. The HADDOCK docking models for the BM5 new cases were generated using predicted interface residues from CPORT^59^ as reported in the BM5 paper^58^. The docking models for ZDock, pyDock and SwarmDock were taken from the work of Moal et al.^31^. In total, we could use 9, 18, 14 and 10 complexes for HADDOCK, SwarmDock, pyDock and ZDock, respectively, with the number of models per case varying from 125 to 500, for which at least one near-native model was present in the set of generated models.

##### Calculating HADDOCK energetic terms

We used HADDOCK to calculate the intermolecular energies for the docking models from other docking programs. For this, the missing atoms of the models were built according to the OPLS force field topology with standard HADDOCK scripts using CNS^60^. A short energy minimization (EM) was then performed with the following settings: 50 steps of conjugate gradient EM, van der Waals interactions truncated below the distance of 0.5Å, and dielectric constant set to 1.

##### Removing docking models containing clashes

Docking models originating from rigid-body docking programs, such as ZDock and pyDock, often contain clashes that a short EM cannot resolve. We removed those clashing models from the test dataset following the CAPRI assessment procedure: A clash is defined by a pair of heavy atoms between protein partners with a distance below 3Å. We discarded all models with more than 0.1 clashes per Å^2^ of buried surface.

##### Clustering

The remaining docking models for each case were clustered with the fraction of common contacts (FCC) method^61^ using a 0.6 cutoff and requiring a minimum number of 4 members per cluster.

#### II. Validation on the CAPRI score set

The CAPRI score set consists of a set of models collected from CAPRI participants and used in the scoring experiment of CAPRI^62^. We tested our scoring functions on this dataset and compared its performance with various scoring functions used in the CAPRI challenge. Docking models with clashes were removed as described above. Both dimers and multimers were considered here. We used 13 cases from the CAPRI score set with number of models ranging between 497 and 1987. Following the CAPRI assessment protocol, we considered only 10 models for assessment. The selection was conducted with simply selecting the top 2 models of the top 5 clusters for each target.

### Availability

The iScore code is freely available from Github: https://github.com/DeepRank/iScore. And the docking models used are available from SBGrid: https://data.sbgrid.org/dataset/XXX (the deposition to SBGrid will be done at revision time).

## RESULTS

### Training and optimisation

We trained a novel scoring function called iScore based on random walk graph kernels (RWGK), embedding protein-protein interface conservation profiles and integrating three intermolecular energy terms (electrostatics, van der Waals and desolvation energies) (see Methods). A subset of the docking benchmark 4 (BM4)^63^ was used for training, consisting of 117 crystal structures of protein-protein complexes and docking models obtained with the ab-initio docking mode of HADDOCK for 63 out of those 117 complexes for which near-native docking models were obtained in the final HADDOCK water refinement stage (referred to as the BM4 HADDOCK dataset).

We first trained a graph kernel-based scoring function called GraphRank using a SVM classifier. GraphRank ranks docking models based on their similarity/dissimilarity to the native/non-native set of structures used in the training. The similarity is measured concerning interface topology and conservation. For this, we represent the interface of a protein-protein complex by a graph, using interface residues as the nodes of the graph and intermolecular residue-residue contacts within 6Å as graph edges. The graph nodes are labelled with values of interface residue conservation profiles from PSSM. A novel RWGK based on the framework of Vishwanathan et al.^37^ was designed to measure the similarity between two interface graphs. It was used to train a SVM model on a balanced dataset consisting of 117 native and non-native structures, respectively. The resulting model or scoring function named GraphRank is then used to rank docking models. It takes as input the graph similarity of a docking model with the 234 structures in the training set. The smaller the GraphRank score is, the more similar the docking model is to native complexes.

We then trained iScore by integrating the GraphRank score with three intermolecular energy terms from HADDOCK (see Methods). iScore consists of a linear combination of those four features whose weights were optimized on the BM4 HADDOCK docking models. To avoid extreme values of energies, we independently normalised the various terms for each complex with their median and interquartile range values. The iScore function with its optimised weights is:

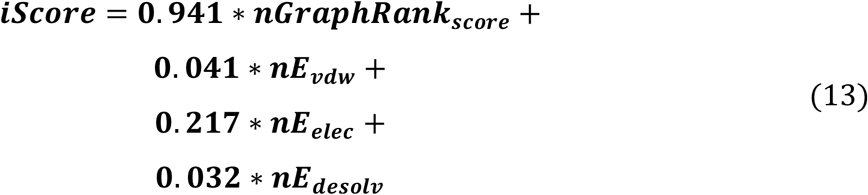

where *nGraphRank_score_, nE*_*vdw*_, *nE*_*elec*_, and *nE_desolv_* are the normalized GraphRank score, Evdw, Eelec and Edesolv energies, respectively.

The success rates of HADDOCK score, GraphRank score and iScore on the BM4 HADDOCK dataset (63 complexes) are shown in Figure 2. Compared with the energy-based HADDOCK score, the graph- and conservation-based GraphRank score has higher success rates. It is also evident that adding energetic features in iScore results in an improved scoring, reaching a success rate of 62% on the top 5 clusters in comparison with 59% for GraphRank.

**Figure 2.**
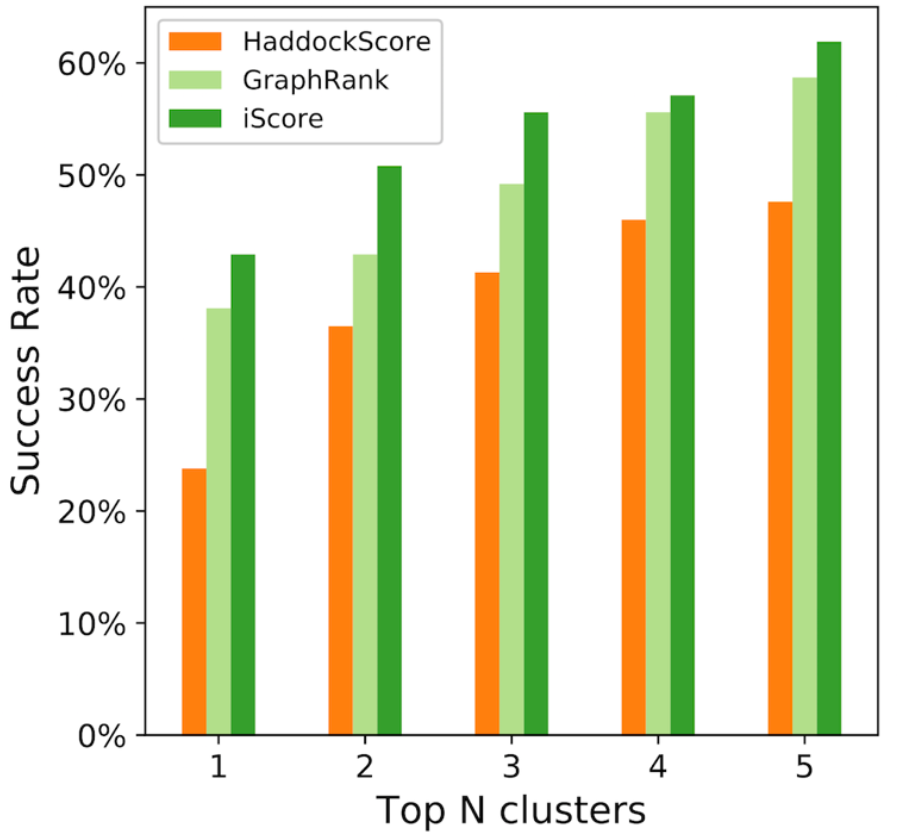
Success rate of HADDOCK score, GraphRank and iScore on the BM4 HADDOCK training dataset over top N clusters of models.

### Benchmarking on docking software-specific docking models and their respective scoring functions

Sampling and scoring are typically not independent components. They are often interrelated since a specific scoring method might depend on the sampling strategy followed and the representation of the system. We benchmark here the performance of iScore and GraphRank (which are trained on HADDOCK models) on docking software-specific docking models and compare their performance with that of each software respective scoring function.

For this, models from the new protein-protein complexes of Docking Benchmark 5^58^ generated using four widely used docking programs: HADDOCK^16,57^, SwarmDock^18^, pyDock^19–21^ and ZDock^22,23^. The number of available complexes with near-native docking models for those four widely-used docking programs are 9, 18, 14 and 10, respectively, with the number of docking models per complex varying from 125 to 500. The scoring performance was assessed with clustering of the docking models using our cluster procedure descried in Methods.

iScore outperforms HADDOCK, ZDOCK and pyDock scoring functions and competes with that of SwarmDock on their respective docking program-specific models (Figure 3). On the HADDOCK models (Figure 3A), iScore shows the same performance as GraphRank, both outperforming HADDOCK on the top2 to top4, reaching 33% success rate for top 5 clusters. For all the other model sets, iScore outperforms GraphRank. It shows a better scoring performance than the original scoring functions of pyDock (Figure 3C) and ZDock (Figure 3D), while the original SwarmDock scoring function remains the best in terms of scoring performance (Figure 3B). iScore reaches a success rate of 36% and 60% (top 5 clusters) on pyDock and ZDock models, respectively, which is clearly a great improvement.

**Figure 3.**
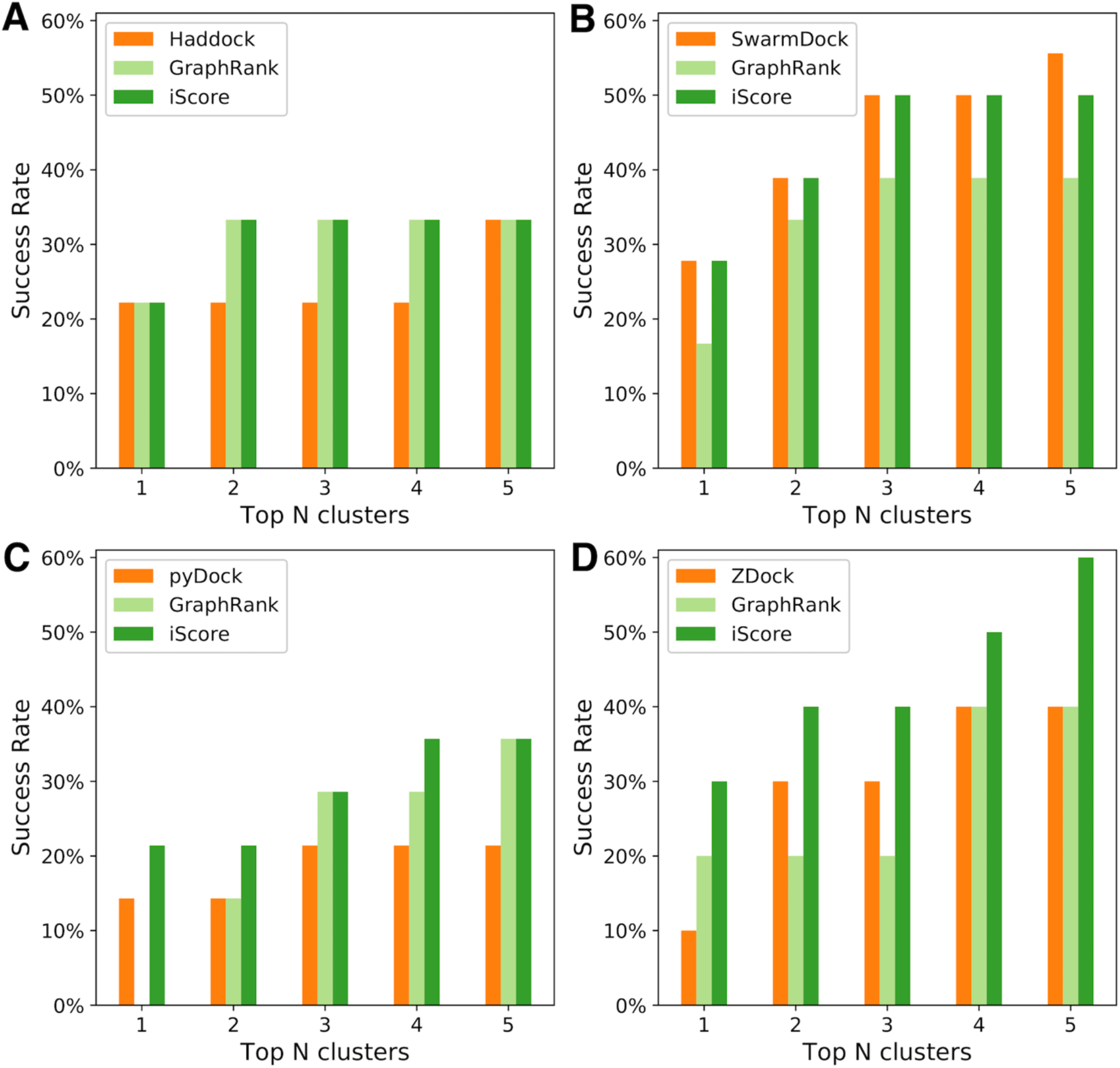
Success rates measured at cluster level on four sets of docking program-specific models for BM5 protein-protein complexes. GraphRank and iScore are compared with scoring functions from HADDOCK (**A**), SwarmDock (**B**), pyDock (**C**) and ZDock (**D**) on the docking models of the corresponding docking program, respectively.

### iScore ranks among the top scorers on the CARPI score set

The scoring set from the CAPRI scoring experiments^62^ is a valuable resource for evaluating scoring functions. CAPRI is a community-wide experiment for evaluating docking programs (started in 2001)^64^ and scoring functions (from 2005 on). The CAPRI score set consists of 15 targets, 13 of which have near-native docking models. Each target has a mixture of 500-2000 models from the various docking programs used in the CAPRI prediction challenges (Table 1). This represents an ideal set for evaluating scoring functions *independently of docking programs*.

**Table 1.**
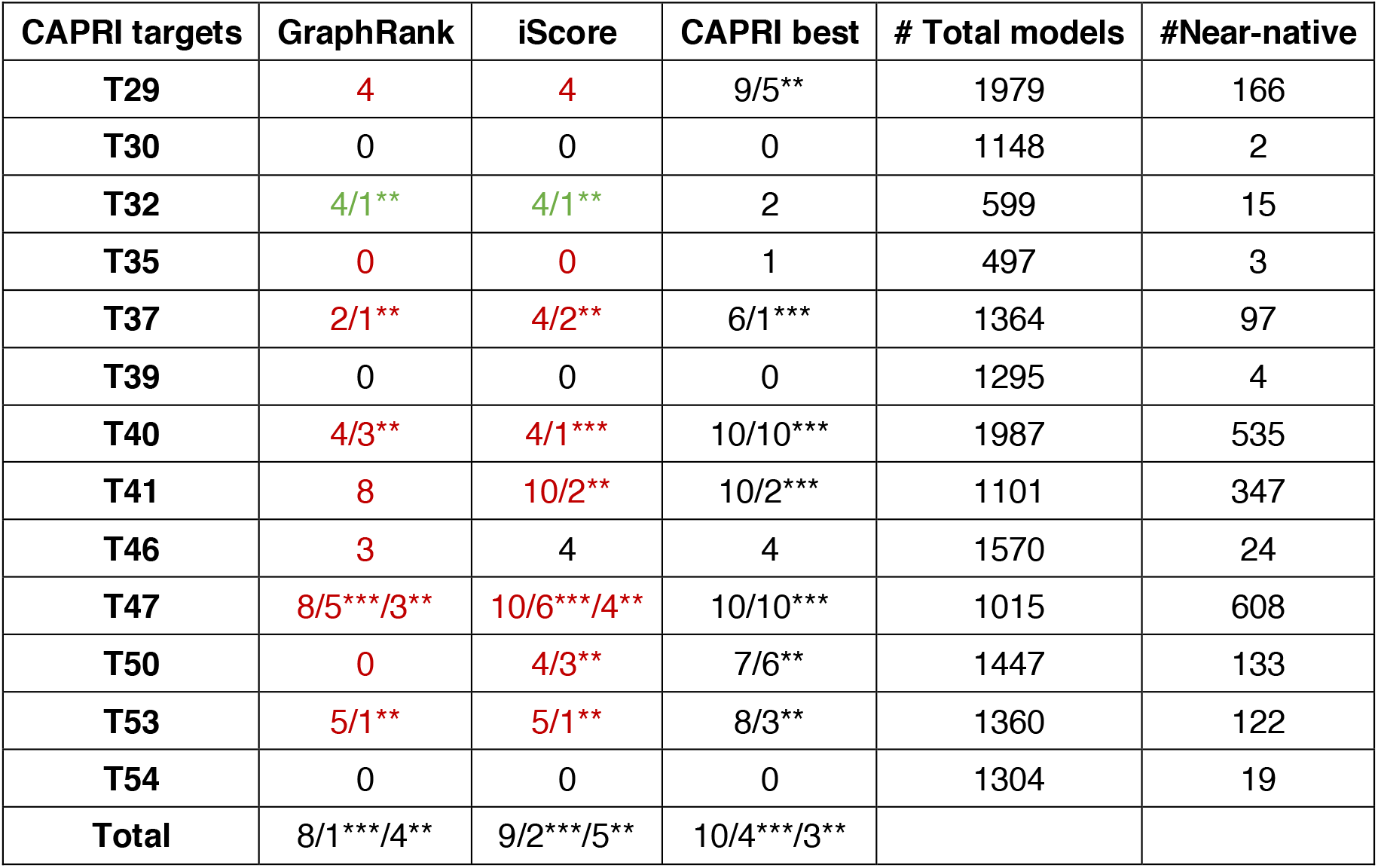
Comparison of GraphRank and iScore with CAPRI best performing group per target on the CAPRI score set. 10 models are selected and evaluated. The values are labelled in green/red when the performance of our scoring functions is better/worse than the CAPRI best scoring group. The scoring performance for each target is reported as the number of acceptable or better models (hits), followed by the number of high (indicated with ***) or medium quality models (**). For example, 8/2** means that there are totally 8 hits among the top 10 models, 2 models out of which are medium-quality models. The overall performance of each method on all 13 targets (the last row) is reported in a similar way. For example, 9/2***/5** means that a scoring function is successful in 9 targets, 2 targets out of 9 have at least a *** model, and 5 out of 9 have at least a ** model in the top 10. Note that the CAPRI best column consists of results from 37 different groups (refer to Table 2 for a comparison of the performance per group and **Table S2** per target).

**Table 2.**
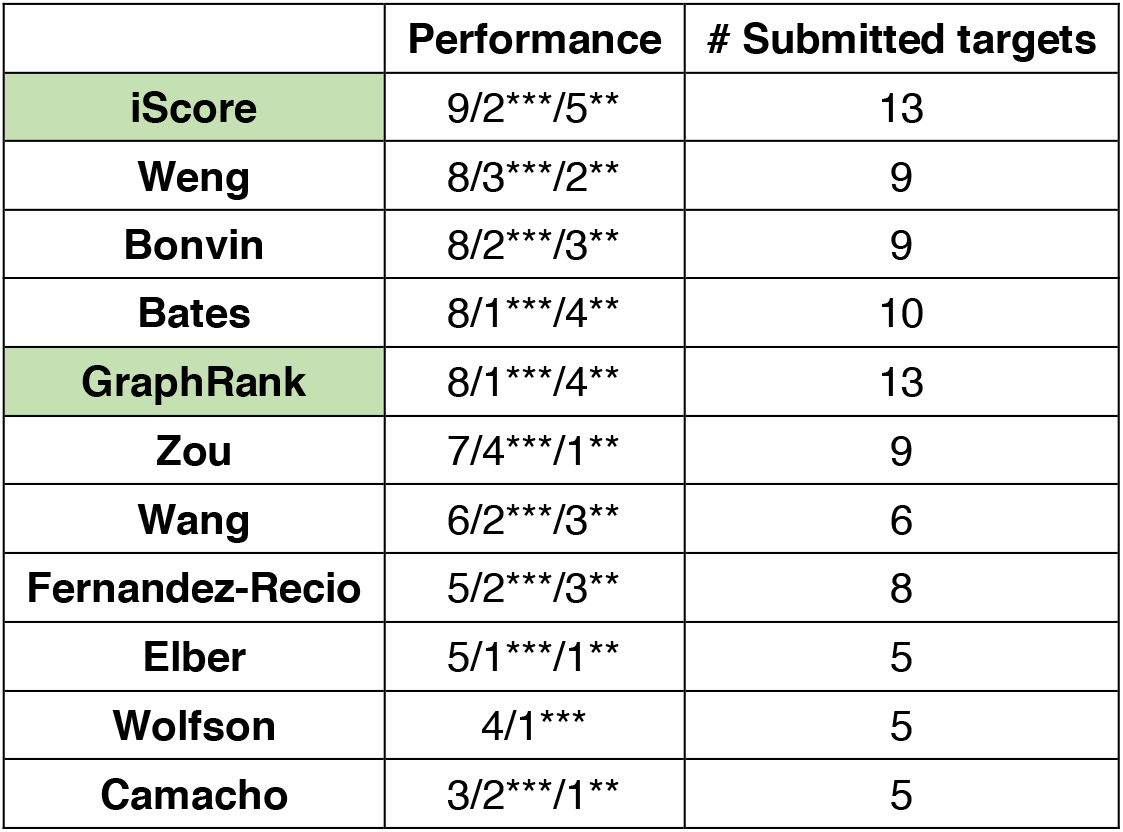
Rankings of GraphRank and iScore in comparison with the scorer groups on the CAPRI score set. In total 37 scorer groups were assessed (**Table S2**), but only scorer groups that have submitted predictions for at least 5 out of the 13 CAPRI targets are shown here. The scoring functions/groups are ordered based on their performance. GraphRank and iScore are highlighted in green. Number of targets with submitted predictions are shown for each function/group.

|  | Performance | # Submitted targets |
| --- | --- | --- |
| iScore | 9/2***/5** | 13 |
| Weng | 8/3***/2** | 9 |
| Bonvin | 8/2***/3** | 9 |
| **Bates** | 8/1***/4** | 10 |
| **GraphRank** | 8/1***/4** | 13 |
| **Zou** | 7/4***/1** | 9 |
| **Wang** | 6/2***/3** | 6 |
| **Fernandez-Recio** | 5/2***/3** | 8 |
| **Elber** | 5/1***/1** | 5 |
| **Wolfson** | 4/1*** | 5 |
| **Camacho** | 3/2***/1** | 5 |

We benchmarked iScore and GraphRank on the models from the CAPRI score set and compared their performance with the reported performance of the various scoring functions/groups which participated to the CAPRI scoring experiments. Following the CAPRI assessment protocol, we selected only the top 10 ranked models for assessing the performance of iScore and GraphRank. This was done by selecting the top 2 models from each of the top 5 clusters for each target.

The scoring performance of iScore and GraphRank on the 13 CAPRI targets containing near-native models is summarised in Table 1, together with the performance of the best scoring function/group in CAPRI for each target. Details of the performance of the various scoring functions compared for these targets are available in **Table S2**. Again, iScore outperforms GraphRank (Table 1) demonstrating the synergistic effects of conservation information and the interacting energies in differentiating near-native models from docking artifacts. Further, iScore selected near-native models on the top10 for 9 out of 13 targets, with 2 targets having high-quality models and 5 having medium-quality models. As a comparison, selecting for each target the best CARPI scoring function/group resulted in 10 out of 13 correctly predicted targets, with 4 and 3 targets having at least one high-quality and medium-quality models, respectively.

Overall, iScore ranks among the top scorers on these 13 CAPRI scoring targets (Table 2). In total 37 scoring functions/groups were assessed (**Table S2**), but only those that participated to at least 5 targets are shown in Table 2. When considering the common submitted targets (**Table S2**), iScore still competes with the Weng group (8/2***/4** vs. 8/3***/2**), the Bonvin group (8/2***/4** vs. 8/2***/3**) and the Bates group (8/2***/4** vs. 8/1***/4**). It should be noted that the CAPRI scoring groups, e.g. Weng and Bonvin groups, selected the 10 models with help of human expertise, while our selections were only generated from iScore and GraphRank without manual selection. Furthermore, considering that GraphRank only uses interface residue conservation profile as feature, it is rather impressive that GraphRank was ranked in the top 4.

## DISCUSSION

We have developed a novel graph-kernel based scoring function, iScore, for scoring and ranking docking models of protein-protein complexes. By benchmarking on docking models from four different docking programs, iScore shows competitive or better success rate than the original scoring functions of those docking programs. Further, validation on CAPRI targets and comparison with CAPRI scorer groups highlights the high performance of iScore, which achieves the top success rate with acceptable or better models selected for 9 out of 13 CAPRI targets. This is quite remarkable considering that a rather small dataset was used for training and that only a single feature was used by GraphRank and 4 features in total by iScore. We can expect to further improve the performance of iScore, by increasing the size of the training set and enriching the node and edge labels of interface graphs.

The usage of graph kernel on labelled graphs in iScore provides a novel way to score docking models. SPIDER^36^ is also a graph-based scoring function but is drastically different from our GraphRank hence also iScore. SPIDER identifies common interface residue patterns (i.e. interfacial graph motifs) in native complexes and rank a docking model by counting the frequency of the interfacial graph motifs. First of all, GraphRank is based on graph kernel functions to calculate the interface similarities between a docking model and the training complexes while SPIDER is based on the frequent graph mining technique to identify interfacial graph motifs. Second, and importantly, the graphs used in SPDIER has only node labels with amino acid identity, while our GraphRank framework can potentially explore not only the properties of individual interface residues with node labels, but also the features of contacts between residues with edge labels. While we have only used node labels in this work (residue conservation profiles), the concept can easily be extended to add labels to the graph edges, for example in the form of residue-residue interaction energies. Third, iScore uses multiscale representations of docked interfaces by combining atom-level energy terms with residue-level graph similarities, which allows to account for both subtle differences in 3D space, interaction topology and residue conservations at the same time.

Both conservation profiles and intermolecular energies are important features for scoring of PPIs. Our scoring function GraphRank, using only conservation profiles of the interface residues as features, already shows a promising scoring performance. Physical energies have been widely used and identified as important features in state-of-the-art scoring functions and are complementary to evolutionary information. Considering the successful applications of intermolecular energies in existing scoring functions, in this work we simply combined three intermolecular energetic terms from HADDOCK with the conservation profiles-based GraphRank score. The resulting scoring function iScore outperforms GraphRank, indicating the significance of considering both evolutionary and energetic information in characterizing PPIs.

When comparing the performance of iScore on models from different docking programs on BM5 new data, we do observe iScore is able to improve the ranking over the original scoring functions for the rigid-body docking programs (pyDock and ZDock), while iScore does not really outperform the flexible docking programs like HADDOCK and SwarmDock which generate more optimised interfaces (Figure 3). This might be related to the structure quality of the docking models. For docking models from flexible docking, their structures are already optimised to release steric clashes, while the rigid-body programs usually do not have such an optimisation step, leading to unnatural interactions (clashes) within structures. To improve the structure quality of the docking models, we did apply a short energy minimization to optimise the structures before calculating intermolecular energies. With higher structure quality, like those coming out of SwarmDock and HADDOCK, the impact of this short minimisation is smaller, and the resulting improvement of iScore versus the original scoring functions is less.

By introducing the labelled graphs and graph kernel in our scoring function iScore, we pave the way for exploring more detailed features in the graph presentation of protein-protein complexes. Natural extensions of this work will be to include edge labels, for example residue-residue interaction energies and co-evolution. Considering graphs are natural representations of biomolecules, this general framework should be useful for other important macromolecular interaction related topics, such as binding affinity predictions, hot-spot predictions, and rational design of protein interfaces.

## ACKNOWLEDGEMENTS

This work was supported in part by the European H2020 e-Infrastructure grant BioExcel (grant no. 675728). CG acknowledges financial support from the China Scholarship Council (grant no. 201406220132). LX acknowledges financial support from by the Netherlands Organisation for Scientific Research (Veni grant 722.014.005) and an Accelerating Scientific Discovery (ASDI) grant from the Netherlands eScience Center (grant no. 027016G04). The work of VH was supported in part by the National Center for Advancing Translational Sciences, National Institutes of Health through the grant UL1 TR000127 and TR002014, by the National Science Foundation, through the grants 1518732, 1640834, and 1636795, the Pennsylvania State University’s Institute for Cyberscience and the Center for Big Data Analytics and Discovery Informatics, the Edward Frymoyer Endowed Professorship in Information Sciences and Technology at Pennsylvania State University and the Sudha Murty Distinguished Visiting Chair in Neurocomputing and Data Science funded by the Pratiksha Trust at the Indian Institute of Science. YG was supported in part by a research assistantship fundedby the Center for Big Data Analytics and Discovery Informatics at Pennsylvania State University. We thank Dr. Iain H. Moal (EBI Hinxton, UK) for providing docking models of SwarmDock, pyDock and ZDock. We thank Dr. Yasser EL-Manzalawy from Penn State University and MSc. Mick Walter from Utrecht University for helpful discussions. The content is solely the responsibility of the authors and does not necessarily represent the official views of the sponsors.

