## Supplementary material for "iScore: A novel graph kernel-based function for scoring protein-protein docking models": Table S1,Table S2.

**Support material**

for

**Table S1.** BLAST parameters.

| Query Length | Substitution Matrix | Gap Open Cost | Gap Extend Cost |
|---|---|---|---|
| <35 | PAM-30 | 9 | 1 |
| 35-50 | PAM-70 | 10 | 1 |
| 50-85 | BLOSUM-80 | 10 | 1 |
| >85 | BLOSUM-62 | 11 | 1 |

**Table S2.** Scoring performance on 13 CAPRI targets.

The table summarizes the performance of scoring functions or scorer groups on the 13 CAPRI targets from the CAPRI score set[1].

For each scoring function or scorer group the number of models of each CAPRI category[2]: high quality (***), medium quality (**), and acceptable (*) are listed. When a scoring function or scorer group submitted models of medium or high quality in addition to acceptable models, the number of acceptable models is not annotated with "*". The scoring performances for CAPRI scorer groups are collected from literatures[3,4]. Cases, where no scoring predictions were submitted, are highlighted in grey, those with successful scoring (in the top10) in green and those with no acceptable predictions in red.

The list of all scorer groups or scoring functions in each table is ordered by the total performance. The number of valid targets with available predictions is shown for each scoring function.

| | T29 | T30 | T32 | T35 | T37 | T39 | T40 | T41 | T46 | T47 | T50 | T53 | T54 | Total Performance | #valid |
|---|---|---|---|---|---|---|---|---|---|---|---|---|---|---|---|
| **iScore** | 4* | 0 | 4/1** | 0 | 4/2** | 0 | 4/1***/3** | 10/2** | 4* | 10/6***/4** | 4/3** | 5/1** | 0 | 9/2***/5** | 13 |
| **Weng** | 3/2** | | | | 2/1*** | 0 | 7/2*** | 4* | 3* | 9/6***/3** | 1* | 3/1** | | 8/3***/2** | 9 |
| **Bonvin** | 9/5** | | | | 2/1** | | 10/2*** | 10* | 2* | 10/9***/1** | 2* | 8/3** | 0 | 8/2***/3** | 9 |
| **Bates** | 4/2** | 0 | | | 6/1*** | | 10/9** | 4* | 2* | 10/10** | 2* | 1** | 0 | 8/1***/4** | 10 |
| **GraphRank** | 4* | 0 | 4/1** | 0 | 2/1** | 0 | 4/3** | 8* | 3* | 8/5***/3** | 0 | 5/1** | 0 | 8/1***/4** | 13 |
| **Zou** | | | 0 | | 4/2*** | | 10/2*** | 10/2*** | 1* | 10/10*** | 2/1** | 1* | 0 | 7/4***/1** | 9 |
| **Wang** | | | | 1* | 6/4** | | 8/1*** | | | 2/2*** | 7/6** | 5/1** | | 6/2***/3** | 6 |
| **Fernandez-Recio** | 5/1*** | | 0 | 0 | | 0 | | 3/2** | | 10/4***/6** | 6/1** | 4/1** | | 5/2***/3** | 8 |
| **Elber** | | | | | | | 8/3*** | 1* | 1* | | 2* | 5/1** | | 5/1***/1** | 5 |
| **Wolfson** | 2* | | 0 | | 1* | | 9/6*** | | 1* | | | | | 4/1***/3* | 5 |
| **Camacho** | 2/1** | | 0 | | 0 | | 10/9*** | | | 10/9***/1** | | | | 3/2***/1** | 5 |
| **Haliloglu** | | | | | 5/4** | | 6/4*** | | | 1/1*** | | | | 3/2***/1** | 3 |
| **Gray** | | | | | | | 0 | | | 9/3***/6** | 4/1** | 1** | | 3/1***/2** | 4 |
| **Kihara** | | | | | | | 0 | 3/1** | 4* | 7/6***/1** | | | | 3/1***/1** | 4 |
| **Grudinin** | | | | | | | | | | 5/3***/2** | 1* | 3/1** | | 3/1***/1** | 3 |
| **Xiao** | | | | | | | | | 1* | | 6/4** | 3* | | 3/1** | 3 |
| **Takeda-Shitaka** | | 0 | | | | | 10*** | 10/2** | | | | | | 2/1***/1** | 3 |
| **Korkin** | | | | | | | | | | 10/3***/7** | 1** | | | 2/1***/1** | 2 |
| **Aze** | | | | | 1*** | | | | | | | | | 1/1*** | 1 |
| **Umeyama** | | | | | | | | | | 8/7***/1** | | | | 1/1*** | 1 |
| **Vajda** | 2/1** | | | | | | | | | | | | | 1/1** | 1 |
| **SAMSON+HEX** | | | | | | | | 10/1** | | | | | | 1/1** | 1 |
| **Liu** | | | | | | | 4/4** | | | | | | | 1/1** | 1 |
| **Vakser** | | | 2* | | | | | | | | | | | 1/1* | 1 |
| **Mitchell** | | | | | 2* | | 0 | | | | | | | 1/1* | 2 |
| **Bajaj** | | | | | | | | | | | 1* | | | 1/1* | 1 |
| **Seok** | | | | | | | | | | | | 1* | | 1/1* | 1 |
| **Poupon** | | | | | | | | | | | | 0 | | 0 | 1 |
| **Ten Eyck** | 0 | | | | | | | | | | | | | 0 | 1 |
| **Zhou** | 0 | | | | | | | | | | | | | 0 | 1 |
| **Smith** | 0 | | | | | | | | | | | | | 0 | 1 |
| **FIREDOCK** | | | | | | | 0 | | | | | | | 0 | 1 |
| **Lee** | | | | | | | 0 | | | | | | | 0 | 1 |
| **Elofsson** | | | | | 0 | | | | | | | | | 0 | 1 |
| **Pal** | | | | | | | | | | | 0 | | | 0 | 1 |
